# Transvaginal ovum retrieval in scimitar horned oryx (*Oryx dammah*) and roan antelope (*Hippotragus equinus*)

**DOI:** 10.64898/2026.04.30.721932

**Authors:** Parker M. Pennington, James D. Gillis, Darya A. Tourzani, Christopher J. Lambert, Thanh Q. Nguyen, Steve Metzler, Scott B. Citino, Matt James, Linda M. Penfold, Jason R. Herrick

## Abstract

Development and use of assisted reproductive technologies (ARTs) in non-domestic species provides novel tools for species conservation. As a first step towards in vitro embryo production, we developed an OPU technique for two antelope species, scimitar horned oryx (*Oryx dammah*) and roan antelope (*Hippotragus equinus*) utilizing a custom-made needle guide and existing OPU equipment utilized by livestock and human practitioners. Females were anesthetized and placed in sternal recumbency for transvaginal OPUs. Prior to OPUs (36 - 45 hours), SHO and roan were either hormonally stimulated with follicle stimulating hormone (FSH, 140 or 250IU) as a single injection or not. A total of 32 and 26 OPUs were completed in SHO (n=10) and roan (n=7), respectively, representing one to four OPUs per animal at monthly intervals. A total of 141 oocytes were recovered from 215 follicles in SHO and 31 oocytes from 58 follicles in roan. FSH dose (250IU) increased (P<0.05) the number of follicles aspirated and the number of oocytes recovered in SHO. No effects of FSH were observed in roan (P>0.05). Good quality oocytes were recovered from all females and procedures were conducted in four consecutive months with no evidence of scar tissue buildup or reduced capacity to recover quality oocytes. These ARTs can be used to develop in vitro embryo production tools for population management and the preservation of female genetics; bolstering genetic diversity and guarding against extinction.

## INTRODUCTION

Development of assisted reproductive techniques (ARTs) for non-domestic species advance our understanding of species biology and provide novel tools for species preservation (Mastromonico, 2024, Herrick, 2019). Similar ARTs developed and utilized in domestic species can serve as a basis for use in non-domestic and endangered counterparts, but differences in both husbandry and biology often require species-specific modifications of domestic animal protocols (Morrow et al., 2009). Researchers have made progress adapting various ARTs for use in endangered ungulate species (Pukazhenthi, 2016), but efficiency remains low compared to domestic species. ARTs in wildlife have focused predominantly on semen collection and cryopreservation and/or artificial insemination, presumably because male gametes are more easily acquired than oocytes. (Morrow, 2009, Pukazhenthi et al., 2006, Penfold et al., 2005; review, Pukazhenthi, 2016). Complimentary protocols to collect oocytes for in vitro embryo technologies are equally important for population management and species conservation but have received far less attention (Mastromonico, 2024).

Previous use of ARTs in nondomestic hoofstock include techniques like artificial insemination, routine reproductive ultrasonography, estrous cycle manipulation, and embryo recovery and transfer (Pope et al., 1991; Schiewe et al., 1991; Wirtu et al., 2009; Morrow et al., 2009; Pukazhenthi, personal communication). Nondomestic species are typically less tractable than domestic counterparts and there may be welfare risks associated with repeated handling. (Pennington et al., 2013). Additionally, differences in estrous cycle parameters between domestic and non-domestic species reduce the efficiency of applying synchronization and stimulation protocols developed for domestic species directly to non-domestic species (Morrow et al., 2009). While ARTs hold promise for endangered species, they have not been utilized to their full potential because resources are limited in traditional zoological institutions. In the meantime, female genetic representation is limited and maintenance of genetic diversity in managed populations is at risk of becoming unsustainable (Mastromonico, 2024).

The development of ARTs for conservation of female genes, such as ovum retrieval, would provide alternative tools to augment population management. There are multiple approaches to ovum retrieval from females, including transvaginal and laparoscopic approaches. The transvaginal route offers a less invasive option for larger species like cattle (Review: Chinarov, 2024), horses (Stout, 2023), and pigs (Oltedal et al., 2024), while the laparoscopic route accommodates ovum collection from smaller species like goats and sheep (Baldassarre, 2021). Both approaches have become commercially viable options in cattle, horses, and small ruminants (Chinarov, 2024; Stout, 2023; Baldassare, 2021), demonstrating their efficacy and utility in domestic species. However, adaptation of these procedures for species like antelope that don’t have a direct domestic model species requires additional optimization due to anatomical differences (i.e. body size, cervical anatomy, or uterine tone). Transvaginal ovum retrieval has been completed in eland (*Taurotragus oryx*), bongo (*Tragelaphus eurycerus*), addax (*Addax nasomaculatus*) and banteng (*Bos javanicus*) (Wirtu et al., 2009; Asa et al., 1998; Halls-Woods, 2000), while the laparoscopic approach has been used in gazelle (*Gazella dama mohrr, Berlinguer* et al., 2008). Prior to this study, OPU had not been attempted in scimitar horned oryx (*Oryx dammah*) or roan (*Hippotragus equinus*) antelope.

While SHO are endangered in the wild following reintroduction efforts thanks to captive breeding efforts, these efforts have also resulted in a robust captive population providing animals for research. As a result, ART protocols have been developed for SHO, including the production of offspring from both AI and ET (in vivo produced embryos) (Pope et al., 1991; Schiewe et al., 1991; Morrow et al., 2009, B. Pukazhenthi, personal communication). While roan are listed as least concern in the wild, populations are declining and are considered endangered in some regions of Africa (IUCN.org). Captive populations of roan are small and mostly based in Africa. A small population has been established in the United States, but it is unlikely to remain sustainable, as has been reported for other populations of antelope (Wildt et al., 2019). Few studies have been conducted on roan antelope regarding reproduction, and all focus on endocrine parameters (Kamgnang et al., 2023) and no attempts have been made to develop ART. In the current study, we describe the development of transvaginal OPU in SHO and roan that could be used to provide oocytes for ART in these and other antelope species.

## METHODS

### Animals

This study was reviewed and approved by Colossal’s Ethics and Welfare in Research Committee (protocol number CEWR25001PP). Female-only herds of scimitar horned oryx (n=10, SHO) and roan (n=7) antelope were maintained at a private, USDA-licensed facility. SHO (92-130 kg) were of mixed parity and roan (127-156.5 kg) were nulliparous and all were sexually mature at the initiation of the study. Animals were housed such that social and species specific behaviors could be maintained throughout the study.

SHO were housed in enclosures (6000sqft) and maintained on perennial peanut hay, concentrate (ADF 16, Mazuri), and water *ad libitum*. Roan were maintained on a 4.5 acre pasture, supplemented with peanut hay and concentrate (ADF 16, Mazuri) and provided water *ad libitum* throughout the study. Oocyte collections on both species were conducted once per month with a minimum of 21 days between procedures.

Procedures were conducted on females of both species at random stages of the estrus cycle and no attempt to synchronize cycles prior to OPU procedures was made. Preliminary studies in each species indicated 2-5 follicles were present on the ovaries of each animal (data not shown) at any given exam. To maximize the potential of the procedures and increase the size and number of follicles present at the time of OPU, females were treated with a single injection of follicle stimulating hormone (FSH,140 or 250 IU/animal), a method that has been previously used in cattle (Vieira et al., 2016) to reduce injection number, animal handling, and stress. Briefly, lyophilized FSH (Folltropin®, 700IU per bottle) was reconstituted in sodium hyaluronate (GEL-50, 5mg/ml) to achieve doses of either 140 or 250 IU/animal, delivered by dart (<2 ml injection volume) 36 – 45 hours prior to OPU procedures. The initial dose (140 IU) and subsequent dose (250 IU) were based on those used for small ruminants (Mendes et al., 2018). The same females were included in both the control (No FSH) and treatment (140 or 250 IU FSH) groups, with specific treatments outlined in Table 1.

**Table 1:** Summary of results from OPU procedures in individual scimitar horned oryx (SHO) and roan for four consecutive months. SHO procedures occurred October 2025 through January 2026 while roan OPUs occurred November 2025 through February 2026. Shaded cells indicate FSH stimulation (250 IU) prior to OPU.

| Species | October |  | November |  | December |  | January |  |
| --- | --- | --- | --- | --- | --- | --- | --- | --- |
| SHO | Oocytes | Follicles | Oocytes | Follicles | Oocytes | Follicles | Oocytes | Follicles |
| 3 | 5 | 10 | 8 | 9 | 9 | 11 |  |  |
| 4 | 2 | 4 | 10 | 10 | 5 | 5 | 15 | 16 |
| 5 | 1 | 2 | 2 | 4 | 1 | 2 | 2 | 5 |
| 7 | 2 | 3 | 4 | 7 | 0 | 1 |  |  |
| 10 | 7 | 9 | 3 | 5 | 2 | 6 |  |  |
| 12 | 0 | 4 | 10 | 10 | 3 | 4 | 5 | 6 |
| 14 | 4 | 8 | 1 | 2 | 2 | 6 |  |  |
| 16 | 4 | 7 | 13 | 20 | 4 | 12 | 4 | 6 |
| 2 |  |  |  |  |  |  | 10 | 13 |
| 40 | 0 | 2 | 1 | 2 | 2 | 4 |  |  |
| Totals | 25 | 49 | 52 | 69 | 28 | 51 | 36 | 46 |
| FSH Dose | 140 IU |  | 250 IU |  | 250 IU |  | 250 IU |  |

|  | November |  | December |  | January |  | February |  |
| --- | --- | --- | --- | --- | --- | --- | --- | --- |
| Roan | Oocytes | Follicles | Oocytes | Follicles | Oocytes | Follicles | Oocytes | Follicles |
| 1 | 2 | 2 | 0 | 2 | 2 | 3 | 1 | 1 |
| 2 | 1 | 2 | 0 | 1 | 1 | 4 | 2 | 1 |
| 3 | 0 | 1 | 1 | 3 | 2 | 2 | 1 | 2 |
| 4 | 0 | 2 | 1 | 2 | 0 | 2 | 0 | 3 |
| 5 | 2 | 4 | 1 | 2 | 1 | 1 | 3 | 3 |
| 6 |  |  | 1 | 2 | 4 | 3 | 1 | 3 |
| 10 |  |  | 2 | 3 | 1 | 1 | 1 | 3 |
| Totals | 5 | 11 | 6 | 15 | 11 | 16 | 9 | 16 |
| FSH Dose |  |  |  |  | 250 IU |  | 250 IU |  |

### Anesthesia

OPU procedures were conducted under general anesthesia administered and monitored by a licensed veterinarian. For SHO, anesthesia was induced with a combination of butorphanol (27.3 mg/mL), azaperone (9.1 mg/mL), and medetomidine (10.9 mg/mL) (BAM, 2mL, Wedgewood Pharmacy) supplemented with ketamine (100 mg) delivered by dart (Pneu-Dart, Type P, 3.0 ml, IM). For roan, anesthesia was induced with a combination of thiafentanil (1.2 – 1.8 mg), xylazine (20-28 mg), and ketamine (100 - 150 mg) delivered by dart (Pneu-Dart, Type P, 1.5 – 2.0 ml, IM). Darts were projected with a Pneu-Dart G2 X-Caliber gas-based rifle. Once recumbent, animals were loaded onto a flatbed trailer and moved to a covered location. In SHO, anesthesia was maintained by supplementation with ketamine (50 – 100mg, IV). In roan anesthesia was maintained by ketamine supplementation (50 – 100 mg, IV) and/or by guaifenesin drip in 5% dextrose (IV). During the procedure, all individuals were supplemented with Cydectin (40 - 50 mg), Multimin (Zinc, Manganese, Selenium, Copper, 3 mL), Oxytetracycline (LA300; 8 - 12 mL) and BO-SE (Selenium, Vitamin E, 4 - 5 mL), and Banamine (100 - 200 mg). Following the completion of each OPU, the animal was returned to either pen or pasture and recovered using a combination of atipamezole (100 mg) and naltrexone (50 mg) for SHO and Naltrexone (50 mg) and yohimbine (30 mg) for roan, given IM. Animals were then monitored until standing. Post-recovery all animals were observed throughout the day and daily thereafter.

### Ovum Pickup

Once recumbent, animals were weighed and a reproductive ultrasound exam was conducted to identify structures on the ovary (corpus luteum, follicle number and size). Briefly, the rectum was voided of feces and an ultrasound probe (MindRay Vetus E7, rectal linear, 7 mHz) was introduced into the rectum with a gloved and lubricated hand to image the ovaries and uterus. Simultaneously, the animal was evaluated for body condition and the condition of their hooves and horns were evaluated. If needed, hooves were trimmed and horns addressed for cracks or breaks.

Once the initial exams were complete, the perineal area was cleaned with soap and water and dried with a clean cloth. The tail was held out of the area for the duration of the ovum retrieval to prevent contamination of the field. The OPU device consisted of an endo-cavity transducer (MindRay V11-3Ws, 4.5 –9.5mHz) with a custom needle guide affixed to the dorsal portion (Figure 1, patent pending). The needle assembly consisted of a 20ga needle affixed to the end of a hollow stainless-steel rod and tubing assembly (WTA, #21789) connecting the OPU needle (WTA, #17925) and collection receptacle (50mL conical tube). The collection receptacle was also connected to the aspiration pump (Cook Medical, #K-MAR-5200-US), with pressure adjusted per species (SHO: 6-7mL/minute, ∼50 mmHg; Roan: 15-16mL/min, ∼100mmHg), creating a closed system. Additionally, all connection points, including the filter to the aspiration pump, were wrapped in parafilm to ensure a tight seal and consistent negative pressure during aspiration. A small amount of warmed (39 °C) and heparinized (10 IU/mL) embryo flush medium (ABT Complete Flush, AgTech) primed the aspiration tubing prior to use for each animal/procedure such that ∼2-5mL of warm flush medium was in the conical tube at the start of each procedure.

**Figure 1.**
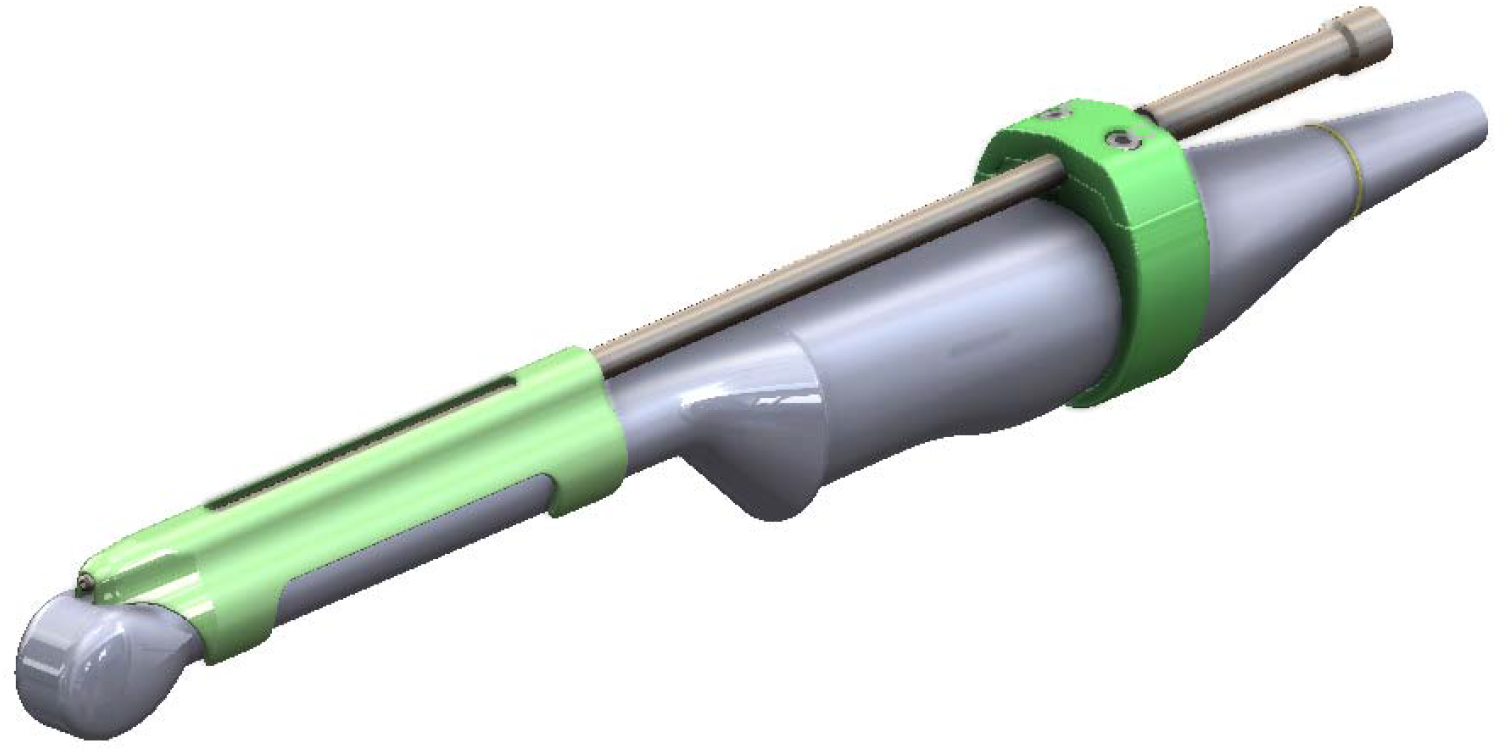
CAD rendering of custom designed, 3D-printed needle guide attachment to ultrasound transducer for ovum pickup. The green portion indicates a 3D printed component (green) securing a commercially available needle guide to the ultrasound transducer.

Sterile lubricant was applied to the imaging portion of the OPU device, which was then covered in a chemise and additional sterile lubricant was applied. The prepared OPU device was introduced to the vagina and seated adjacent to the cervix. The receptacle was maintained in a tube warmer (WTA, #24390). A gloved and lubricated arm was then introduced to the rectum and each ovary was digitally manipulated per rectum to the vaginal wall adjacent to the OPU device for imaging (Figure 2). The needle assembly was introduced to the needle guide and the aspiration pump was turned on. Follicles were aligned with the projected needle path indicated on the ultrasound screen and the needle was advanced into the follicle and contents were aspirated (Figure 2). The needle was rotated ∼180 degrees in each direction before withdrawing fully into the needle guide. Once all visible and accessible follicles were aspirated, all instruments were removed from the animal.

**Figure 2:**
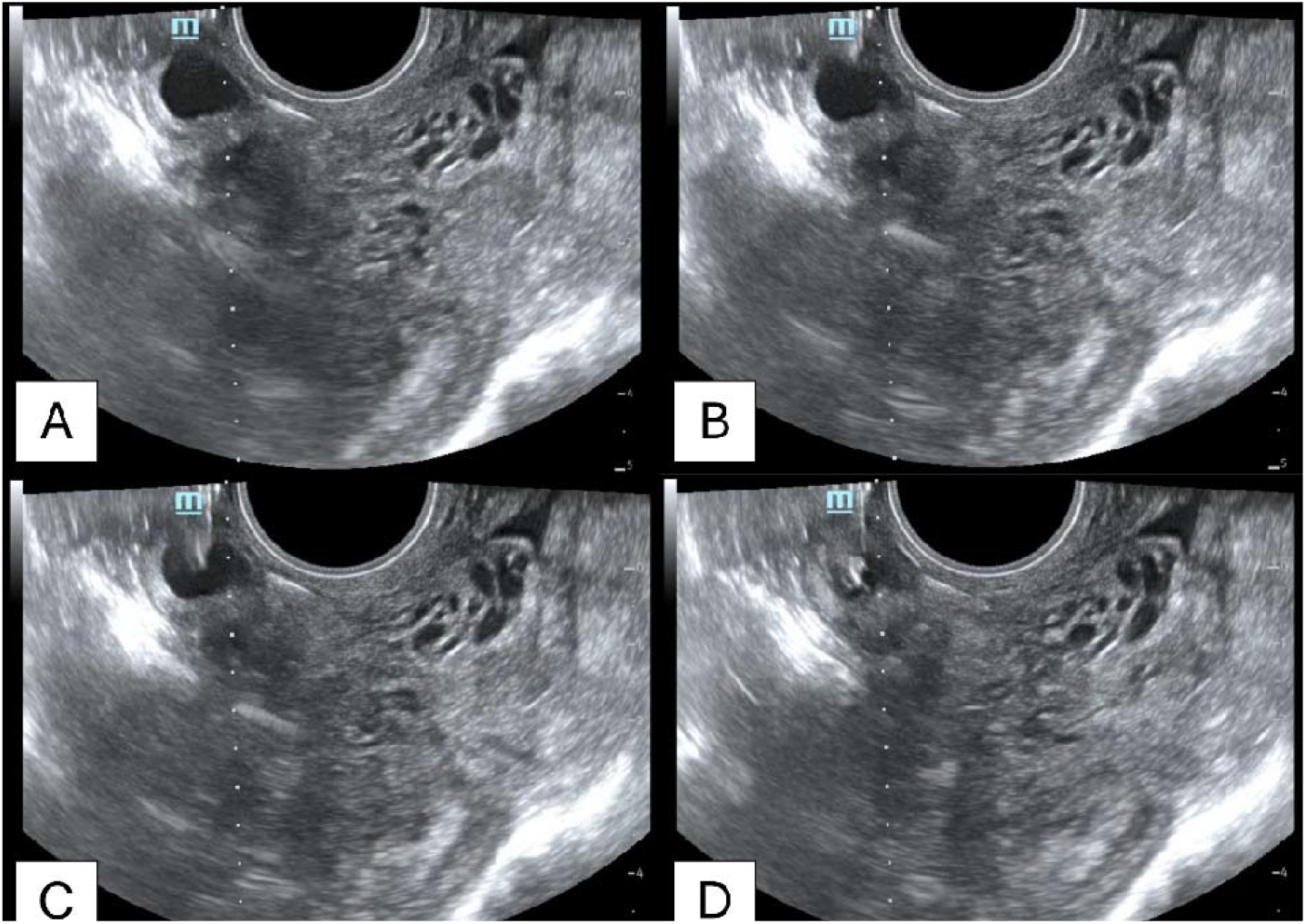
Representative ultrasound images of ovarian follicle alignment, needle puncture, and collapse. Images depict (A) alignment of the ovarian follicle with the needle trajectory indicated by the dotted white line, (B) introduction of the needle through the vaginal wall and into the field of view, (C) puncture of the follicle with the needle, and (D) collapse of the follicle through aspiration at 50 mmHg of pressure. The white arrow indicates the needle entering the visual field and follicle.

Between animals, the OPU apparatus was disassembled and cold sterilized in chlorhexidine (0.2%), rinsed in distilled water, then rinsed in sterile saline before being reassembled. A new needle assembly was used for each animal.

### Oocyte Recovery

Once the OPU procedure was complete, the needle assembly was flushed with warm, heparinized medium to ensure any oocytes remaining in the line were recovered. The receptacle was then disconnected from the aspiration tubing and rinsed. The aspirate was then taken to a clean space and searched for oocytes using a stereomicroscope with a heated stage (39°C). Recovered oocytes were washed through clean media (x3) before being placed in maturation medium (IVF Bioscience) for subsequent studies of in vitro maturation and fertilization.

### Statistics

A one-way ANOVA was used to analyze the differences between the number of follicles aspirated and the number of oocytes recovered following 0, 140, and 250 IU FSH treatments. Tukey’s HSD post-hoc test was used to determine differences between groups in the case of significant ANOVA findings. In roan, a t-test was used to analyze differences between the number of follicles aspirated and oocytes recovered following treatment with 0 or 250 IU FSH. Differences were considered significant when P-values were <0.05. Means are presented ± standard error of the mean.

## RESULTS

Ovum pickup procedures were conducted for four consecutive months, between October 2025 and January 2026 in SHO and November 2025 to February 2026 in roan, each month representing one OPU attempt per animal (Table 1). The average procedure time (initial immobilization dart to animal recovered and standing) was 1 hour for both species. No adverse side effects have been observed following any of the procedures. Similarly, no signs of infection or internal injury were identified in any individuals during the study based on visual observations and ultrasound imaging of the reproductive tract.

In SHO, a total of 32 OPUs was conducted in 10 females (up to 4 procedures per animal) (Table 1, 2), while a total of 26 OPUs was conducted in 7 roan females (up to 4 OPUs per animal) (Table 2). Prior to initiation of the study, hymen were manually ruptured in roan in preparation for OPU procedures. Hymen were mildly vascularized and appeared to partially reform, evidenced by the need to break down remnant membranes at subsequent procedures. Two roan females joined the study group in the second month, increasing the number of animals in the study to 7. However, they were not treated with FSH prior to OPUs to reduce potential stress during their first procedures.

**Table 2:** Oocyte recovery rate in unstimulated (No FSH) and stimulated (FSH) scimitar horned oryx (SHO) and roan. A significant difference was found between groups in follicles aspirated (P=0.004), and oocytes recovered (P=0.01) in SHO. No significant differences were found (P>0.05) between No FSH and FSH groups in either follicles aspirated or oocytes recovered in roan. All values reported as mean ± SEM.

| Species<br>(n=females) | Hormone<br>Stimulation | Procedures | Follicles<br>Aspirated | Oocytes<br>Recovered | Oocyte Recovery<br>Per Follicle (%) |
| --- | --- | --- | --- | --- | --- |
| **SHO**<br>**(n=10)** | No FSH | 14 | 4.0 ± 0.6<sup>a</sup> | 2.4 ± 0.6<sup>c</sup> | 49.6 ± 8.3% |
| | 140 IU FSH | 4 | 7.5 ± 1.6<sup>a,b</sup> | 4.5 ± 1.0<sup>c,d</sup> | 61.0 ± 6.6% |
| | 250 IU FSH | 14 | 9.2 ± 1.3<sup>b</sup> | 6.4 ± 1.2<sup>d</sup> | 64.5 ± 6.6% |
|  | **Total** | **32** | **215** | **141** | **57.6%** |
| **Roan**<br>**(n=7)** | No FSH | 15 | 2.2 ± 0.9 | 1.1 ± 0.3 | 47.4 ± 10.2% |
| | 250 IU FSH | 11 | 2.3 ± 0.3 | 1.3 ± 0.3 | 70.3 ± 17.6% |
|  | **Total** | **26** | **58** | **31** | **57.2%** |

Oocytes were successfully recovered from all individuals in both species (Figure 3). A total of 141 oocytes were recovered from approximately 215 follicles in SHO across 32 total OPU procedures (Table 2). We were able to aspirate a range of follicle sizes (0.2 – 2.1cm) and routinely recover good quality oocytes (Class I – IV, Representative Figure 3). The mean (± SEM) number of oocytes collected per female was 4.0 ± 0.6, 7.5 ± 1.6, and 9.2 ±1.3 from females treated with 0, 140, and 250 IU of FSH, respectively; representing recovery rates (oocytes per follicle) of 49.6 ± 8.3%, 61.0 ± 6.6%, and 64.5 ± 6.6% (Table 2). A significant difference was observed in the number of follicles aspirated (P=0.002) and the number of oocytes recovered (P=0.01) when SHO females received 250 IU FSH, compared to females that did not receive FSH. Good quality oocytes were used for in vitro maturation studies (data not shown).

**Figure 3.**
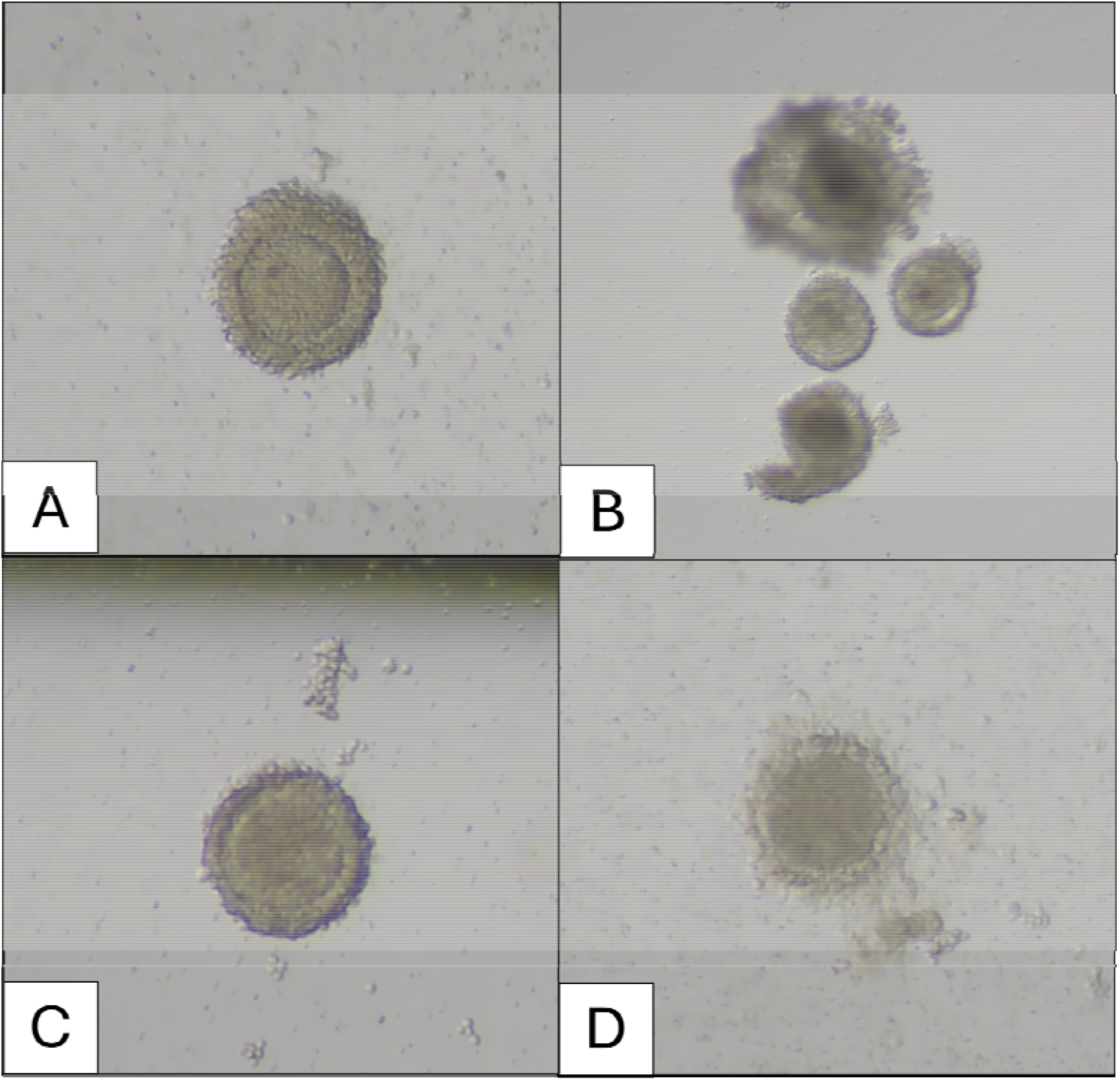
Representative images of oocytes from SHO (A, B) and roan (C, D) at recovery. SHO oocytes (A, B) were recovered under ∼50mmHg of pressure (∼6-7mL/min flow rate) and roan oocytes (C, D) were recovered under ∼100mHg of pressure (∼15-16 mL/min flow rate).

In roan, a total of 31 oocytes were recovered from approximately 58 follicles across 26 total OPUs. We were able to aspirate a range of follicle sizes (0.2 – 1.5cm) and routinely recover good quality oocytes (Class I – IV, Representative Figure 3). Recovery rates were 47.4 ± 10.2% and 70.5 ± 17.6% from non-FSH and FSH stimulated (250IU) OPUs, respectively. Mean oocyte recovery per female was 1.1 ± 0.3 and 1.3 ± 0.3 from non-FSH and FSH stimulated OPUs, respectively (Table 2). Treatment with FSH did not affect (P>0.05) the number of follicles present at the time of OPU or the number of recovered oocytes. Recovered oocytes were utilized in in vitro maturation studies (data not shown).

## DISCUSSION

We report the first successful, repeatable transvaginal ovum pickup in two antelope species: scimitar horned oryx and roan antelope. Although OPU has been utilized in other domestic species, its use in non-domestic species is limited (Wirtu et al., 2009, Woods-Hall, 2000, K. Bauman, personal communication). When oocytes from wildlife have been available for study, they are most often from post-mortem samples in gamete rescue attempts (de Oliviera Santos et al., 2022). Here we show that the technique commonly used in domestic cattle and horses can be adapted and applied to species like SHO and roan through adaptation of the OPU apparatus itself. To accommodate the smaller frame and reproductive tract of these antelope species, we designed a custom needle guide attachment for an ultrasound transducer originally designed for human vaginal use, which provides a wide image array in a small housing. This modification allowed a non-surgical, minimally invasive approach, in contrast to the laparoscopic OPU procedure often employed in domestic small ruminants like goats and sheep (Baldassare, 2021). Interestingly, even though the roan are larger antelope, the same custom probe could be used in both species, although this is likely due to the young age of the individuals used in this study. Commercial OPU often use a microconvex probe and some even offer a ‘heifer’ probe to accommodate the smaller frame of heifers, but these would still be too large for SHO. Similarly, Wirtu et al., (2009) developed an OPU probe with reduced size to accommodate eland – the largest antelope species. It seems that, despite a similar body size of some nondomestic ungulates to domestic cattle, vaginal and rectal sizes remain smaller, and therefore need species-specific consideration.

We performed OPUs on a monthly basis and found it to be safe and effective in SHO and roan. No negative effects of the repeated OPU procedures were observed on ovarian morphology or function in either species based on ultrasound examination of the reproductive tract and consistent oocyte yield. Similarly, food intake and BCS remained stable throughout the study, as did herd dynamics and daily behaviors observed by animal care staff. No adverse impacts of FSH treatment were observed. However, individual variation and response still existed as the number of follicles varied considerably from month to month.

Specialized handling systems, such as chute systems and hydraulic restraint devices, would facilitate routine handling and repeated hormone injections and would likely increase oocyte recovery. However, these systems are not available at most zoological institutions. Therefore, we utilized a single injection protocol for hormonal stimulation to increase the number and size of follicles available for aspiration. An initial dose of 140IU per animal was utilized in SHO based on small ruminant doses for OPU (Mendes et al., 2018), but it did not seem to have any effect on follicles aspirated (P>0.05). The dose was then increased to 250IU per animal, in order to account for the increased weight and body size of SHO compared to small ruminants. This increased dose had a significant impact on the number of follicles aspirated and oocytes recovered (compared to no FSH treatment) in SHO. Subsequent studies with increased sample sizes should focus on combining estrous synchronization with FSH stimulation to fully maximize the potential of OPU. An FSH dose was successfully delivered as a single injection in roan, but no significant differences were found in either follicles aspirated or oocyte recovery. As the roan are larger than the SHO, it is plausible that a larger dose of FSH may be required to achieve the same ovarian response. Additionally, basic studies to characterize the follicular dynamics of both SHO and roan would greatly aid these endeavors (Mantano et al., 2022; Adams et al., 1992), provided that care is taken to minimize stress with the associated increase in handling needed for the monitoring. Despite variability in follicle numbers, the procedure we have developed has immediate and broad applicability to virtually any institution and could even be used for free-ranging wildlife, where animals have not been, or cannot be, treated prior to collection.

All procedures were conducted under a surgical plane of anesthesia with at least 21 days between procedures. To maximize the efficacy and efficiency of these immobilizations, the overall health of each animal was evaluated during each procedure, including weight, body condition, and hoof and horn confirmation. This process allowed us to identify and treat minor issues that are difficult to detect in animals maintained on pasture until overt clinical manifestations are visible. For example, minor hoof issues were found in several of the animals before noticeable changes in their gait had occurred. These issues were treated and resolved quickly (within 2-3 months) due to the routine access to the animals. Similarly, regular assessment of body weight allowed for more precise dosing of anesthetic drugs and detection of subtle changes in weight before a significant change in BCS had occurred.

The repeatability, safety, and efficacy of OPU in SHO, roan, and other medium-sized antelope enables routine and systematic study of female gametes. With improved access to oocytes, advanced embryo technologies like in vitro fertilization, somatic cell nuclear transfer, stem cell line derivation (Mastromonaco, 2024; Pukazhenthi et al., 2006), and other techniques yet to be realized become potential tools for wildlife conservation. Furthermore, oocyte collection provides an opportunity to preserve female genes, either by direct cryopreservation of oocytes or cryopreservation of embryos produced by in vitro fertilization. With these technologies, cryopreserved oocytes and embryos can be integrated with ongoing efforts to cryopreserved spermatozoa, providing a ‘secondary insurance population’ – one that takes up very little space and contains as much of the genetic diversity from the current population as possible.

We show, for the first time, transvaginal OPU is effective in two species of antelope, the SHO and roan. Success was achieved through development of a custom needle guide affixed to an ultrasound probe smaller than that used with commercial transvaginal OPU in domestic species. Treatment with a single injection of FSH prior to OPU (36-45 hrs) increased the numbers of both follicles aspirated and oocytes recovered, providing an efficient and impactful protocol, even without estrus synchronization. The combination of these elements (single FSH injection, no synchronization) provides valuable opportunities to develop additional ART tools for endangered wildlife while maintaining low(er) stress and good wellbeing. Further refinement of protocols for FSH treatment, possibly in combination with estrous cycle synchronization, would be needed to increase the number of oocytes recovered. Through these modifications we were able to achieve a minimally invasive method of oocyte retrieval, expanding the avenues for advanced ARTs that rely on in vitro embryo development, as well as oocyte and embryo banking. This procedure can be expanded to other medium sized antelope species, furthering its potential use as a tool for population management and species conservation.

